# Mitochondrial-Y chromosome epistasis in *Drosophila melanogaster*

**DOI:** 10.1101/807768

**Authors:** J. Arvid Ågren, Manisha Munasinghe, Andrew G. Clark

**Author notes:** These authors contributed equally (J. Arvid Ågren, Manisha Munasinghe).

## Abstract

The coordination between mitochondrial and nuclear genes is crucial to eukaryotic organisms. Predicting the nature of these epistatic interactions can be difficult because of the transmission asymmetry of the genes involved. While autosomes and X-linked genes are transmitted through both sexes, genes on the Y chromosome and in the mitochondrial genome are uniparentally transmitted through males and females respectively. Here, we generate 36 otherwise isogenic *Drosophila melanogaster* strains differing only in the geographical origin of their mitochondrial genome and Y chromosome to experimentally examine the effects of the uniparentally inherited parts of the genome, as well as their interaction, in males. We assay longevity and gene expression through RNA-sequencing. We detect an important role for both mitochondrial and Y-linked genes, as well as extensive mitochondrial-Y chromosome epistasis, in both traits. In particular, genes involved in male reproduction appear to be especially sensitive. Despite these interactions, we find no evidence that the mitochondrial genome and Y chromosome are co-adapted within a geographic region. Overall, our study demonstrates a key role for the uniparentally inherited parts of the genome for male biology, but also that mito-nuclear interactions are complex and not easily predicted from simple transmission asymmetries.

## Introduction

The co-evolution between mitochondrial and nuclear genes is one of the oldest and best studied examples of symbiosis [1–3]. Orchestrated interaction between genes in the two genomes is essential for a number of eukaryotic traits, especially metabolism and energy production [4–6], and this intimate coordination has been taken as evidence for positive selection for cooperative mito-nuclear combinations [7, 8]. Moreover, there has also been a well-documented transfer of genes from the mitochondrial to the nuclear genome [9–11], and animal mitochondrial genomes contain only 37 genes. Finally, the case for the importance of adaptive mito-nuclear epistasis is further strengthened by the observation that placing mitochondrial genomes on novel nuclear backgrounds is often, though not always, associated with adverse fitness effect (see for example Reinhardt et al. [12] and Eyre-Walker [13] for alternative perspectives).

A key factor governing mito-nuclear co-evolution is the difference in transmission between the two genomes. For example, because the mitochondrial genome is almost exclusively maternally inherited [14], mutations that are deleterious in males can spread in a population, given that they are beneficial or neutral in females [15–20]. The occurrence of male-deleterious mitochondrial mutations is particularly well studied in plants [21–24], where such mutations usually prevent pollen production in hermaphroditic plants, essentially rendering individuals female, guaranteeing the mutation’s transmission through ovules. This phenomenon is therefore called cytoplasmic male sterility. In the zoological literature, following Gemmell et al. [15], the presence of mitochondrial mutations with deleterious effects in males is known as the Mother’s Curse.

Other than the mitochondrial genome, other uniparentally inherited genes are those located on the sex-determining chromosome, i.e. on the Y in an XY system where males are the heterogametic sex and on the W in a ZW female heterogametic system. The strict paternal inheritance makes the Y an especially interesting candidate for mito-nuclear epistasis. In particular, it has been suggested to be an ideal location for modifiers that counteract the male deleterious effects of mitochondrial mutations [25, 26], a scenario formally modelled by Ågren, Munasinghe and Clark [27]. Despite its heterochromatic structure and paucity of protein coding genes, the Y chromosome is now recognized as being able to affect a wide variety of traits [28–30]. For example, in *Drosophila* it underlies variation in traits ranging from susceptibility to bacterial infection [31], male reproductive success [32], to sex-specific aging [33].

The extent of mito-Y interactions and their importance to male fitness remain poorly understood. Some suggestive evidence comes from empirical work by Innocenti et al. [34], who found that loci in the mitochondrial genome can affect the expression of a large number of autosomal loci in male, but not in female, *Drosophila melanogaster*. Such sexual dimorphism in expression is consistent with a sex-specific selective sieve being central to mitochondrial genome evolution. Furthermore, several of the loci identified by Innocenti et al. [34] to be sensitive to mitochondrial variation overlap with loci shown by Lemos et al. [29] to be sensitive to variation on the Y chromosome. The extent to which male autosomal gene expression is subject to mito-Y interactions, however, is unclear. If they are important, one prediction may be that males that carry a Y-chromosome and a mitochondrial genome that have co-evolved in the same sympatric population may differ from males where the mitochondrial genome and Y chromosome are from diverged populations and therefore represent a novel mito-Y combination.

Recent attempts to empirically address these questions in the fruitfly *D. melanogaster* have revealed some suggestive patterns. Yee et al. [25] used combinations from three populations (a total of 9 mito-Y combinations) to provide a proof-of-concept evidence of how mito-Y combinations may affect aspects of male fitness. However, they did not find evidence that males with sympatric mito-Y combinations had higher fitness than males with novel combinations. Similarly, Dean et al. [26] found that both mitochondrial and Y-linked genes independently affected male locomotor activity, but only under certain diets and social environments.

Here, we extend these studies in three ways. First, we increase the sample size considerably, by including 36 mito-Y combinations of *D. melanogaster* males with mitochondrial and Y chromosomes sampled from five worldwide locations (Table 1). Second, we assay longevity, another major fitness trait previously shown to be sensitive to mito-nuclear epistasis [35–38]. Finally, we performed RNA-sequencing on all lines to identify the importance of mitochondrial and Y chromosome variation, as well as mito-Y epistasis, for differential gene expression. In line with Dean et al. (2015) and Yee et al. (2015), we found an important role for both mitochondrial and Y-linked genes and an abundance of mito-Y chromosome epistasis in both longevity and autosomal gene expression.

**Table 1.** Geographical origin of strains used to generate mito-Y combinations. Sympatric combinations are shown on white background and novel on grey.

|  |  | Y chromosome genotype |  |  |  |  |  |
| --- | --- | --- | --- | --- | --- | --- | --- |
| Mitochondrial genotype |  | **B04**<br>**Beijing** | **B11**<br>**Beijing** | **N03**<br>**Netherlands** | **N07**<br>**Netherlands** | **ZWH23**<br>**Zimbabwe** | **ZW139**<br>**Zimbabwe** |
|  | **B38**<br>**Beijing** | Beijing<br>×<br>Beijing | Beijing<br>×<br>Beijing | Beijing<br>×<br>Netherlands | Beijing<br>×<br>Netherlands | Beijing<br>×<br>Zimbabwe | Beijing<br>×<br>Zimbabwe |
|  | **B39**<br>**Beijing** | Beijing<br>×<br>Beijing | Beijing<br>×<br>Beijing | Beijing<br>×<br>Netherlands | Beijing<br>×<br>Netherlands | Beijing<br>×<br>Zimbabwe | Beijing<br>×<br>Zimbabwe |
|  | **I02**<br>**Ithaca** | Ithaca<br>×<br>Beijing | Ithaca<br>×<br>Beijing | Ithaca<br>×<br>Netherlands | Ithaca<br>×<br>Netherlands | Ithaca<br>×<br>Zimbabwe | Ithaca<br>×<br>Zimbabwe |
|  | **N01**<br>**Netherlands** | Netherlands<br>×<br>Beijing | Netherlands<br>×<br>Beijing | Netherlands<br>×<br>Netherlands | Netherlands<br>×<br>Netherlands | Netherlands<br>×<br>Zimbabwe | Netherlands<br>×<br>Zimbabwe |
|  | **N02**<br>**Netherlands** | Netherlands<br>×<br>Beijing | Netherlands<br>×<br>Beijing | Netherlands<br>×<br>Netherlands | Netherlands<br>×<br>Netherlands | Netherlands<br>×<br>Zimbabwe | Netherlands<br>×<br>Zimbabwe |
|  | **N23**<br>**Netherlands** | Netherlands<br>×<br>Beijing | Netherlands<br>×<br>Beijing | Netherlands<br>×<br>Netherlands | Netherlands<br>×<br>Netherlands | Netherlands<br>×<br>Zimbabwe | Netherlands<br>×<br>Zimbabwe |

## Methods

### Drosophila melanogaster strains

Both the longevity and the gene expression assays were performed on isogenic *D. melanogaster* males differing only in the geographical origin of their mitochondrial genome and their Y chromosome (referred to as mito-Y combinations throughout). We used a 6×6 crossing design (Table 1), crossing females from six mitochondrial replacement lines with males from six Y chromosome replacement lines, resulting in male offspring with 36 mito-Y combinations. Mitochondrial genomes came from Beijing, China (2 lines; B38 and B39), Ithaca, NY, USA (1 line; I02), and the Netherlands (3 lines; N01, N02, and N23), and the Y chromosomes from Beijing, China (2 lines; B04 and B11), the Netherlands (2 lines; N03 and N07), and Zimbabwe (2 lines; ZWH123 and YZW139). These populations represent deeply divergent mitochondrial and Y chromosome clades and all lines were chosen from the Global Diversity Lines [39]. If the mtDNA and Y chromosome were from the same geographic region, they are labelled “sympatric” and otherwise they are labelled “novel.”

### Longevity assay

#### Scoring survival

Individual life span was quantified for all 36 mito-Y combinations. Flies were collected upon eclosion, sexed, and placed in vials with approximately 20 male flies in each. 6 replicate vials were used for every mito-Y combination. Every other day, flies were transferred without using CO_2_ to vials with fresh food, and individual deaths were recorded. Investigators scoring deaths were blind to which mito-Y combination a given vial contained. Vials were kept in climate controlled growth chambers at 25 °C, and at a 12:12 hours light:dark cycle. In total around 4,500 individuals were scored (Table S1; Figure S1).

### Differential gene expression

#### RNA extraction and sequencing

3-5 day old males from each of the 36 mito-Y combinations were maintained in vials on standard medium in climate controlled growth chambers with a 12:12 hours light:dark cycle at 25 °C. For each mito-Y combination, 10 males were flash frozen. We used two biological replicates for each combination, with individuals for each replicate being collected from separate crosses performed on the same day.

RNA was extracted from the 10 pooled whole-fly males and RNA seq libraries were prepared using the Lexogen QuantSeq 3’ mRNA-Seq Library Prep Kit FWD. Samples were sequenced in a single-end 75 bp run on a NextSeq500 at the Genomics Facility in the Cornell Biotechnology Resource Center.

#### Read Processing and Alignment

The quality of the RNA sequences was assessed using FastQC (http://www.bioinformatics.bbsrc.ac.uk/projects/fastqc). Trimmomatic was then used to clip adapters, the leading 10 bp, and reads once the average quality of a sliding window of 4 bp dropped below 20 [40]. Reads shorter than 20 bp were dropped from subsequent analysis. Next, reads were aligned to the *D. melanogaster* reference genome (Release 6; [41]) using STAR [42] and HTseq-count was used to determine the raw number of read counts per gene [43].

#### Quantifying gene expression

Differential transcript abundance was analysed using DESeq2 [44]. Read counts were normalized using DESeq2’s internal normalization function estimateSizeFactors, which corrects for both library size and RNA composition bias. Lowly expressed genes were removed from subsequent analysis, such that a given gene was only kept in the dataset if it had at least 20 normalized counts in at least half of the samples. After filtering, 9,533 out of 17,324 (∼55%) of the originally identified genes were kept for subsequent analysis.

To identify nuclear genes sensitive to variation in the mitochondrial genome, the Y chromosome, and mito-Y epistasis, we conducted three separate differential expression analyses using linear models. Because principal component analyses suggested the presence of batch effects (Figure S2), all models were controlled for this. Additionally, surrogate variables were identified and incorporated into all models using SVA to adjust for noise and unmodeled artefacts ([45]; see Figure S2 for details). We characterized a gene as being differentially expressed if they maintain significance after performing independent filtering for multiple test corrections using the Benjamini-Hochberg method set with a false discovery rate of 0.05.

We performed a gene enrichment analysis to determine whether certain gene families were differentially expressed across Y haplotypes, mitochondrial haplogroups, and mitochondrial:Y interactions. We used the R Bioconductor package ‘goseq’ [46] to perform gene ontology (GO) analyses. Taking length bias into account, we identified GO categories as either significantly over- or under-represented using a 0.05 false discovery rate cutoff. REVIGO was used to semantically cluster the lists of enriched GO terms to find a representative subset that could be easily analysed and visualized [47].

To determine whether any differentially expressed genes showed testis- or accessory gland-biased expression, we downloaded data from FlyAtlas, which measures the expression levels of a gene in each adult male [48]. The bias metric used is simply the expression of the gene in the tissue of interest divided by the sum of expression over all other tissues. We considered a gene biased for expression in a tissue of interest if the bias metric for that tissue is greater than 0.5, as that indicates that more than half the reads collected for that gene come from that tissue.

All scripts used for the RNA-sequencing analysis are on GitHub (https://github.com/mam737/mitoY_RNASEQ).

## Results and Discussion

### Extensive variation in lifespan across mito-Y combinations

We detected extensive variation in average lifespan across the 36 mito-Y combinations (Figure 1; Table S1). The independent and epistatic contributions by the Y chromosome and mitochondrial background can be captured by the linear model: *lifespan = mitochondrial type + Y chromosome + mito*Y interaction* (Table 2). This analysis reveals a role for mitochondrial-Y chromosome epistasis in governing longevity (F = 1.805, *P* = 0.00836). However, we find no evidence that individuals with a sympatric mito-Y combination live longer than individuals with a novel combination (Wilcoxon rank sum test, W = 1536800, *P =* 0.5114; Figure 1B).

**Figure 1.**
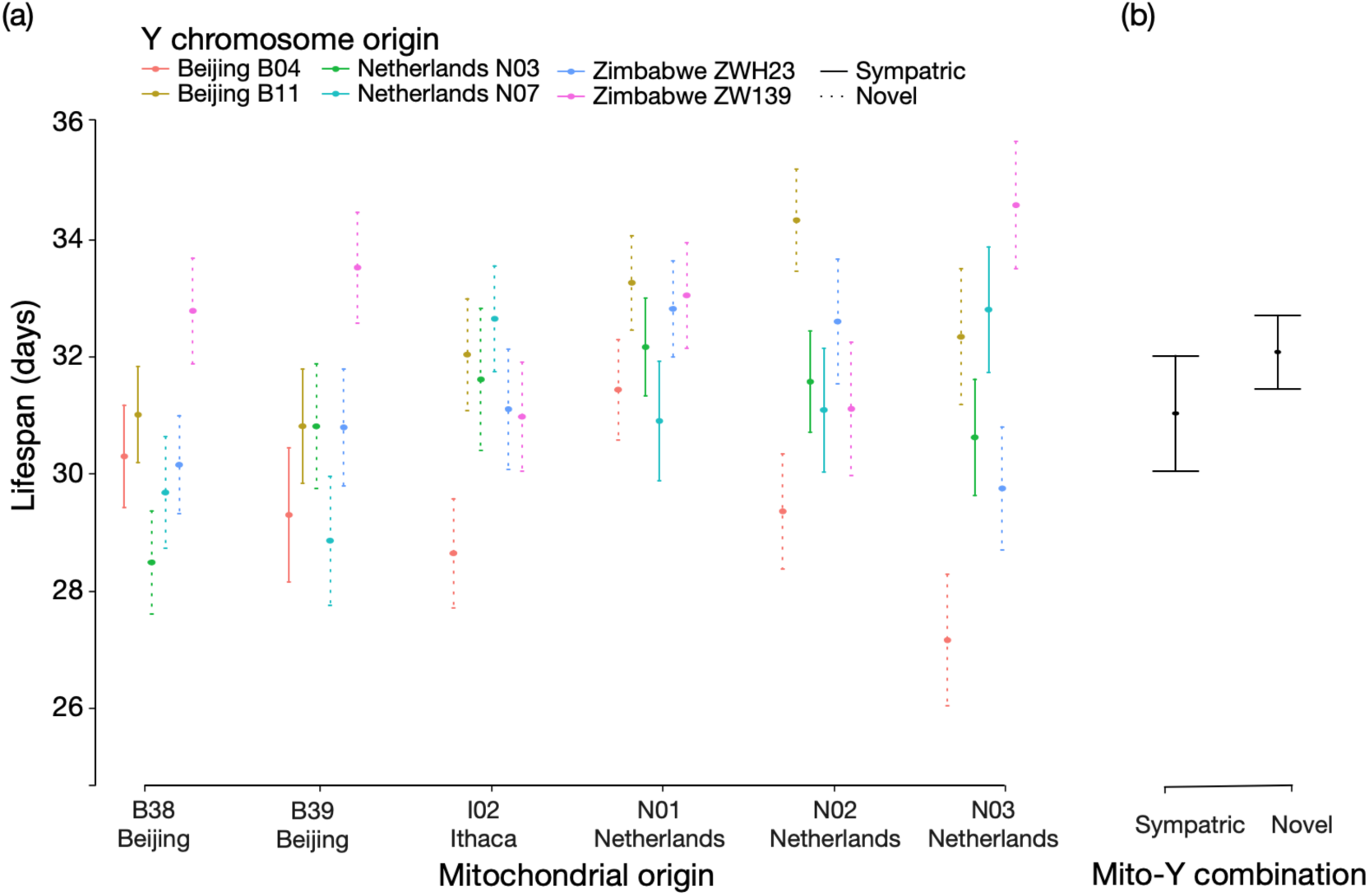
(a) Mean lifespan (days) ± standard error across 36 mito-Y combinations. Geographical origin of the mitochondrial genome is stated at the bottom and colour coded for the Y chromosome. Solid and dashed lines represent sympatric and novel mito-Y combinations respectively. (b) Mean lifespan (days) ± standard error for sympatric (N = 10) and novel (N = 26) mito-Y combinations.

**Table 2.** ANOVA for mito-Y interactions in longevity

| Source | Degrees of freedom | Sum of square | Mean square | F | *P* |
| --- | --- | --- | --- | --- | --- |
| Mitochondrial genome | 5 | 1589 | 317.7 | 3.192 | 0.00703 |
| Y chromosome | 5 | 4000 | 800.1 | 8.039 | $1.49 \times 10^{-7}$ |
| Mito-Y interaction | 25 | 4492 | 1.805 | 1.805 | 0.00836 |
| Residuals | 3789 | 37705 |  |  |  |

To assess how mortality changed over time we fitted a number of survival functions based on different assumptions (non-parametric log-rank, Cox proportional hazard, Gompertz, Gompertz-Makeham, Logistic, and Logistic-Makeham models; see Supplement for details). This comparison revealed significant variation across lines, but no difference between novel and sympatric mito-Y combinations.

As in most species, male *Drosophila* live shorter than females [49, 50], and recent studies have demonstrated a central role for both the mitochondria [51] and the Y chromosome [33] in explaining this difference. We too find that both the Y chromosome and mitochondria, as well as their epistatic interaction, contribute to variation in longevity. Consistent with previous work on mito-Y interactions [25, 26], our results also reveal no evidence that males with sympatric mito-Y combinations differed in these traits compared to males with their mitochondrial genome and Y chromosome sampled from different populations.

### Expression of many nuclear genes is sensitive to variation on the Y chromosome, in the mitochondrial genome or both

We detected 71, 760, and 29 genes whose expression was sensitive to variation on the Y chromosome, in the mitochondrial genome, and mito-Y epistasis respectively. To gain insight into the biological function of these genes, we searched for gene ontology (GO) terms [52, 53] that were either over- or under-represented among our significant hits. We found that genes associated with visual perception (GO:0007601; *P_adj_* = 0.006), response to stimulus (GO:0050896; *P_adj_* = 0.0144), and rhabdomere, a central compartment in compound eyes (GO:0016028; *P_adj_* = 0.0435) were over-represented among genes sensitive to Y haplotype. For those sensitive to mitochondrial haplotype, we find 44 over- and 3 under-represented GO categories with an enrichment of terms belonging to categories such as purine biosynthesis, metabolism, and immune responses (Figure 2). Whereas genes sensitive to mito-Y interactions show substantial variation in expression across samples (Figure 4), we detect no enrichment or depletion of GO categories among the genes sensitive to mito-Y epistasis. Below, we discuss certain biological trends that emerge among our most significant hits.

**Figure 2.**
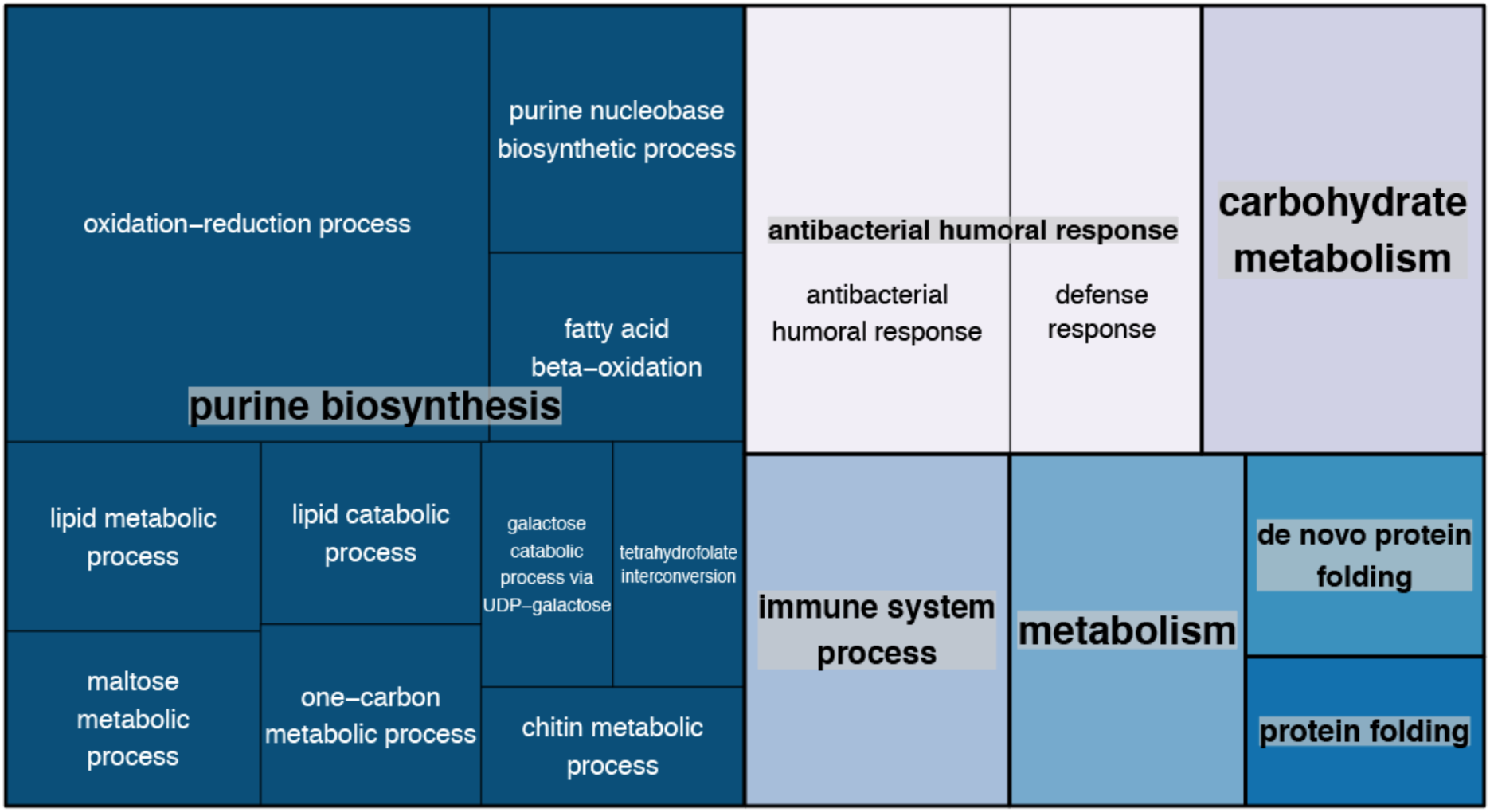
Semantic clustering of over-represented biological process GO terms found among genes sensitive to mitochondrial haplotype. Each rectangle represents a single cluster, which in are joined into larger ‘superclusters’ as visualized by colour. Size of the rectangles indicates *p*-value significance (absolute value of the log 10 transformed *p*-value).

**Figure 4.**
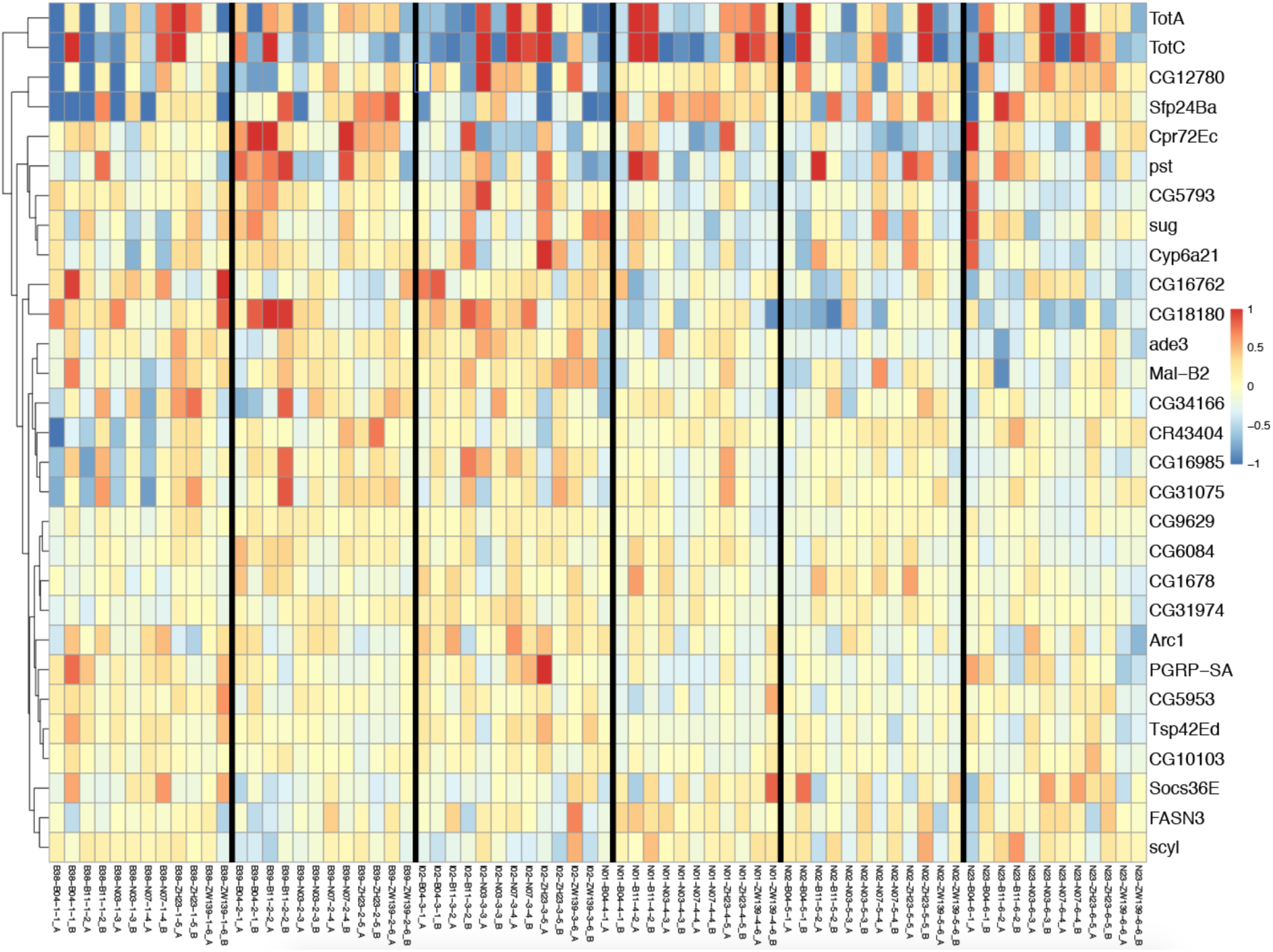
Expression profile of genes sensitive to mitochondrial-Y interactions. Samples (36 mito-Y combinations with 2 replicates, labelled A and B, for each) and genes of interest are listed along the *x*- and *y*-axis respectively. Colour indicates either overexpression (red) or underexpression (blue) compared to the mean expression level of that gene across all samples.

#### Mitochondrial haplotype influences the expression of metabolic genes

Figure 2 highlights the abundance of genes related to metabolism that show differential expression across mitochondrial haplotypes. Many metabolic reactions involve mitochondria, so while the emergence of genes involved in several metabolic processes showing differential expression may not be surprising, the sheer number of nuclear genes whose expression is affected by mitochondrial variation is. Some of these genes exhibit consistent differences between the haplogroups. Several maltases involved in maltose/carbohydrate metabolism show reduced expression among the Netherlands haplogroups compared to the others, whereas 3 of the 4 enzyme-encoding genes involved in the reduction of NADP to NADPH (*Men, Idh,* and *Zw)* show reduced expression in the Beijing B38 haplogroup (Figure S3).

#### Genes related to male fertility show sensitivity to both Y and mitochondrial haplotype

Both the Y chromosome, which contains six essential male fertility factors, and the mitochondrial genome have previously been demonstrated to affect male fertility in *D. melanogaster* [25,32,54–59]. Whereas there is no statistical enrichment of differentially expressed genes belonging to male fertility processes, there are several notable hits that not only show sensitivity across both Y and mitochondrial haplotypes, but also show elevated expression in both the testis and the accessory glands (Figures S5-S8). Among our Y sensitive hits, genes such as *Testis EndoG-Like 1 (Tengl1)*, Adenosine deaminase-related growth factor A2 (*Adgf-A2), Male-specific RNA 98CA (Mst98Ca), gonadal (gdl),* and *Ductus ejaculatorius peptide 99B (Dup99B)* show markedly higher, if not exclusive, expression in either the testis or the accessory glands. Furthermore, *Adgf-A2*, *Mst98Ca,* and *gdl* are all thought to be involved with spermiogenesis, spermatogenesis, or sperm function, while *Dup99B* is a sex-peptide that influences the female post-mating response [60–64].

Several genes sensitive to mitochondrial haplogroup also show higher expression in the testis and accessory glands. Whereas many of these genes are not functionally characterized, there are a few worth highlighting. *Protamine B* (*ProtB)* packages the paternal genome in sperm during spermiogenesis [65, 66], *Otefin (Ote)* encodes a nuclear membrane-associated protein involved in transcriptional silencing of *bag-of-marbles (bam)*, which is a key protein involved in gametogenesis. *Tim17a1*, a subunit of the TIM23 complex, is involved in transporting proteins across the inner mitochondrial membrane and shows markedly higher expression in the testis compared to all other tissues [67]. *Seminase (Sems),* which is expressed predominantly in the accessory glands, is not only transferred during mating but is also thought to be involved in sperm release from storage in females [68]. Finally, we also found several accessory gland proteins, *lectin-46Ca, Acp53C14a*, *Acp33A, CG14034,* and *CG9029. lectin-46-Ca, Acp53C14a,* and *CG13309* (which is significant, but not enriched in the accessory gland) all show reduced expression among the Netherlands mitochondrial haplogroups compared to the others. In addition to genes that are enriched for expression in either the testis or the accessory gland, we find a handful of other genes related to male fertility. *mt:COII* and *mt:Cyt-B* have not only been demonstrated to influence male fertility, but also show no effect on females, leading authors to cite them as examples of Mother’s Curse variants in *Drosophila* [59,69–71]. For *mt:COII*, we see variable expression across mitochondrial haplogroups with the Netherlands N23 haplogroup showing the highest levels of expression (Figure 3b). Another top mitochondrial hit, the heat shock protein *Hsp83*, has been demonstrated to affect spermatid development and differentiation in *D. melanogaster* [72, 73].

**Figure 3.**
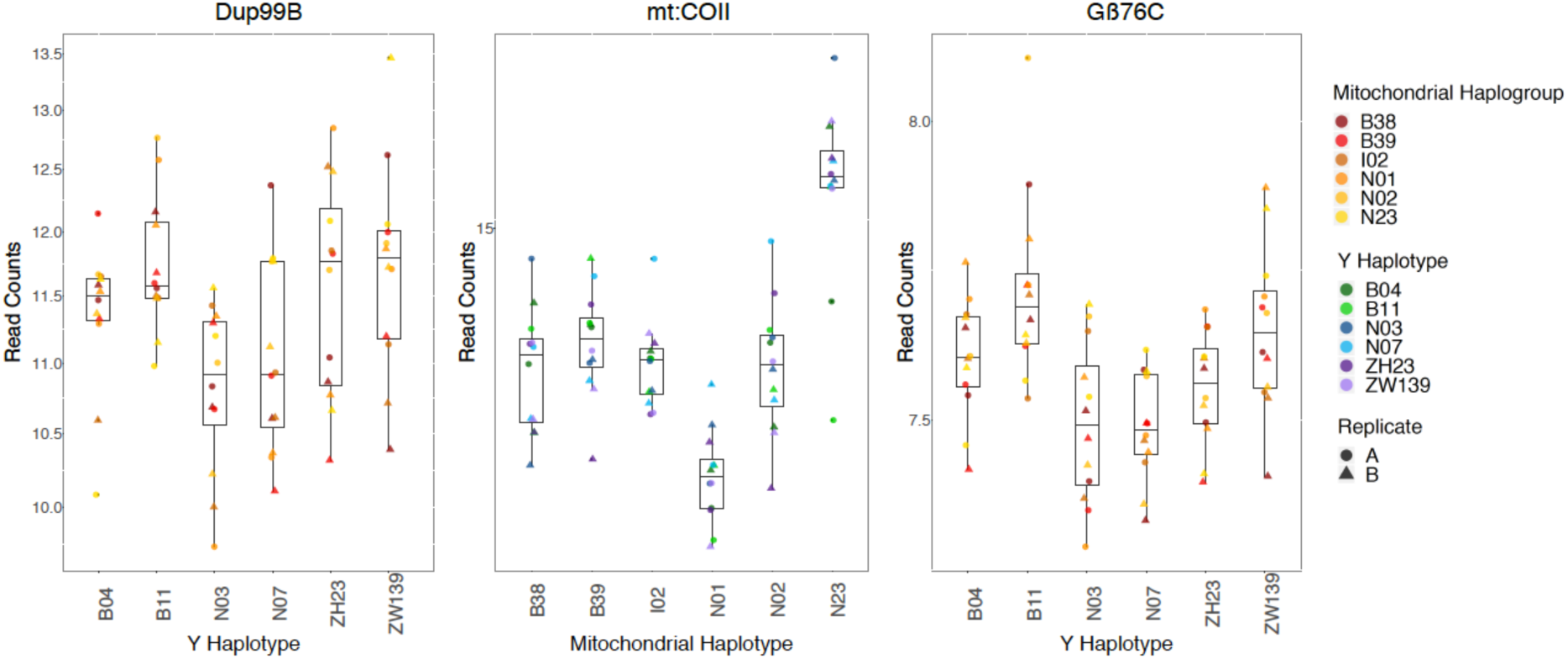
Box and scatter plots visualizing differential expression of selected genes. Normalized read counts for (a) *Dup99B*, a male sex peptide, which shows reduced expression in the Netherlands haplotypes, (b) *mt:COII*, a mitochondrial subunit of cytochrome oxidase II, which shows variable expression across all 6 mitochondrial haplotypes, (c) *Gβ6C*, a Y-sensitive hit, linked to visual perception that shows significant differences in expression across all 6 Y haplotypes.

More surprisingly, we also see differential expression across both Y and mitochondrial haplotypes in genes related to piRNA synthesis. The germline-specific Piwi-interacting RNA (piRNA) pathway protects the genome against transposable elements and viruses and is a highly conserved genomic defence system [74–77]. It restricts transposable element activity by combining the effector function of the Argonaute protein and the specificity provided by the piRNAs [78]. In testes, the piRNA pathway directs this repression through Aub and Piwi, with Piwi playing a vital role in male fertility [75, 79]. While *piwi* and *aub* did not pass our filtering criteria and were subsequently excluded from our analysis, we found that *mino* showed significant differential expression across both Y and mitochondrial haplotypes. *mino* has been previously identified as being critical for primary piRNA biogenesis and is also known to localize to the outer mitochondrial membrane [80]. Two of our mitochondrial sensitive hits, *Hsp83* and *Hop*, are thought to regulate the piRNA pathway through *piwi* to mediate canalization [81].

Despite several genes related to male fertility showing sensitivity to both Y and mitochondrial haplotype, we see little evidence for epistatic interactions between them. Simply looking at the overlap of genes that show sensitivity to Y or mitochondrial haplotype yields only one gene: *mino.* Furthermore, when specifically testing for mito-Y epistasis, the only gene related to male fertility with significant differential expression is the seminal fluid protein *Sfp24Ba*, which, surprisingly, is not found to be significant when testing for sensitivity to just Y or mitochondrial haplotype.

In summary, this analysis identifies several differentially expressed genes that present a strong potential to have downstream consequences for male fertility. The pattern of expression levels seen in Figure 4 suggests no simple additive role of mitochondrial and Y variants, but instead points to specific genotypic combinations having most aberrant expression.

#### Visual perception and nervous system processes show Y haplotype sensitivity

Genes belonging to both visual perception (GO:0007601) and rhabdomere, a key compartment in compound eyes (GO:0016028) are not only enriched among our Y sensitive hits, but they are also some of our most significant hits. We see less expression of these genes among the Netherlands haplotypes, yet markedly higher expression of these genes for the Beijing B11 haplotype (Figure S4). Previous work has shown that variable expression in *Arr1*, *Arr2, Rh4, Gβ6C*, *trpl*, and *Gγ30A* may lead to phenotypic differences in rhodopsin inactivation, phototransduction, and the photoresponse [82–89].

## Conclusions

In this study we used 36 mito-Y combinations to experimentally examine the effects of the uniparentally inherited parts of the genome on male *Drosophila melanogaster*. We detect an important role for both mitochondrial and Y-linked genes, as well as extensive mitochondrial-Y chromosome epistasis in longevity, but little sign that individuals with their Y chromosome and mitochondrial genome sampled from the same population being any different than individuals where the two are from geographically isolated populations. We also detect many genes that are sensitive to variation on the Y chromosome, in the mitochondrial genome or both. The biological function of these genes range from metabolic to visual and neuronal phenotypes, with the strongest effect being for genes involved in male reproduction. Although the results presented here fall short of demonstrating a clear co-evolutionary process between the Y chromosome and mitochondrial genome, the extensive gene expression and longevity interactions highlight the opportunity for these uniparentally inherited segments of the genome to influence male biology.

## Author contributions

JAÅ, MM, and AGC conceived the study. JAÅ performed the fly work and analysed the longevity data. MM analysed the expression data. JAÅ wrote the first draft and all authors contributed to the writing of the manuscript. AGC supervised the project.

## Data and Resource sharing

The RNA-seq data reported here are posted on the GEO resource with reference number xxxxxx. Fly lines and other resources are available on request to AGC.

## Acknowledgements

We thank Amanda Manfredo for help with lab work and Yasir Ahmed-Braimah for guidance on the RNA-sequencing analysis.

## Supplementary Materials

### Analyzing longevity data

To assess how mortality changed over time we fitted several survival functions, under various assumptions (Table S1). A non-parametric log-rank test for survival ∼ mitochondrial-Y combination finds significant difference across lines, but no difference between novel and sympatric combinations (χ^2^ = 124 on 35 degrees of freedom, *P* = 7.81 × 10^-12^; Novel vs. sympatric not significant, *P* = 0.536). We also tested three Cox proportional hazard models, accounting for the mitochondrial genotype (AIC = 55456.55), mitochondrial genotype and Y chromosome (AIC = 55431.19), and finally mitochondrial genotype, Y chromosome, and mito-Y combination (AIC = 55416.34). Comparing AIC scores shows that the last model that includes all three factors provides the best fit. As the name suggests, the Cox model assumes that the risk of dying is constant over time. Testing this assumption reveals it to be violated (*P* = 0.02139), suggesting that age-specific mortality models are more appropriate. A number of such models exist and we consider four of them (the Gompertz, Gompertz-Makeham, the Logistic, and Logistic-Makeham model respectively) using Deday (https://sourceforge.net/projects/deday/). The Gompertz model assumes an exponential increase in mortality with age with α = initial mortality rate (intercept) and β = age-dependent increase in mortality (slope) and the Gompertz-Makeham model is the Gompertz model with an age-independent mortality (*c*).The Logistic model is the Gompertz model with a frailty parameter (*s*), which captures late-life mortality deceleration. Finally, the Logistic-Makeham model includes all four parameters (α, β, *c*, and *s*). We fitted all models across all 36 mito-Y combinations and a comparison of AIC scores suggests that the Gompertz model is best for (almost) all crosses. Because the models are nested we can also use likelihood ratio test to compare model parameters. This comparison reveals little difference across the 36 lines (Table S1; Figure S1).

**Figure S1.**
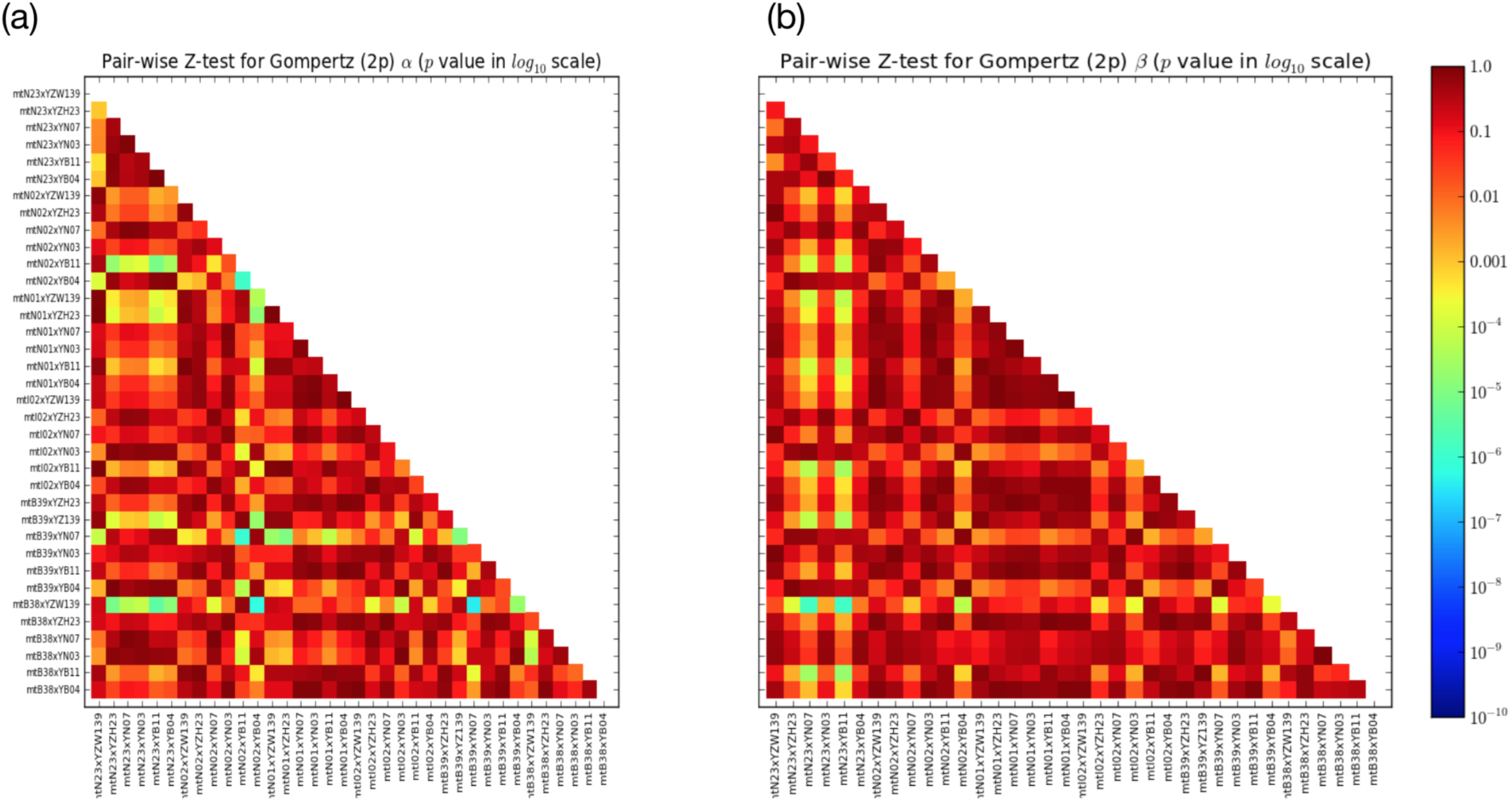
Pairwise Z-tests of α, initial mortality rate (a), and β, age-dependent increase in mortality (b) in the Gompertz model. Reddish colours suggest low significance in difference. Key for genotype names is provided in Table S1.

**Figure S2.**
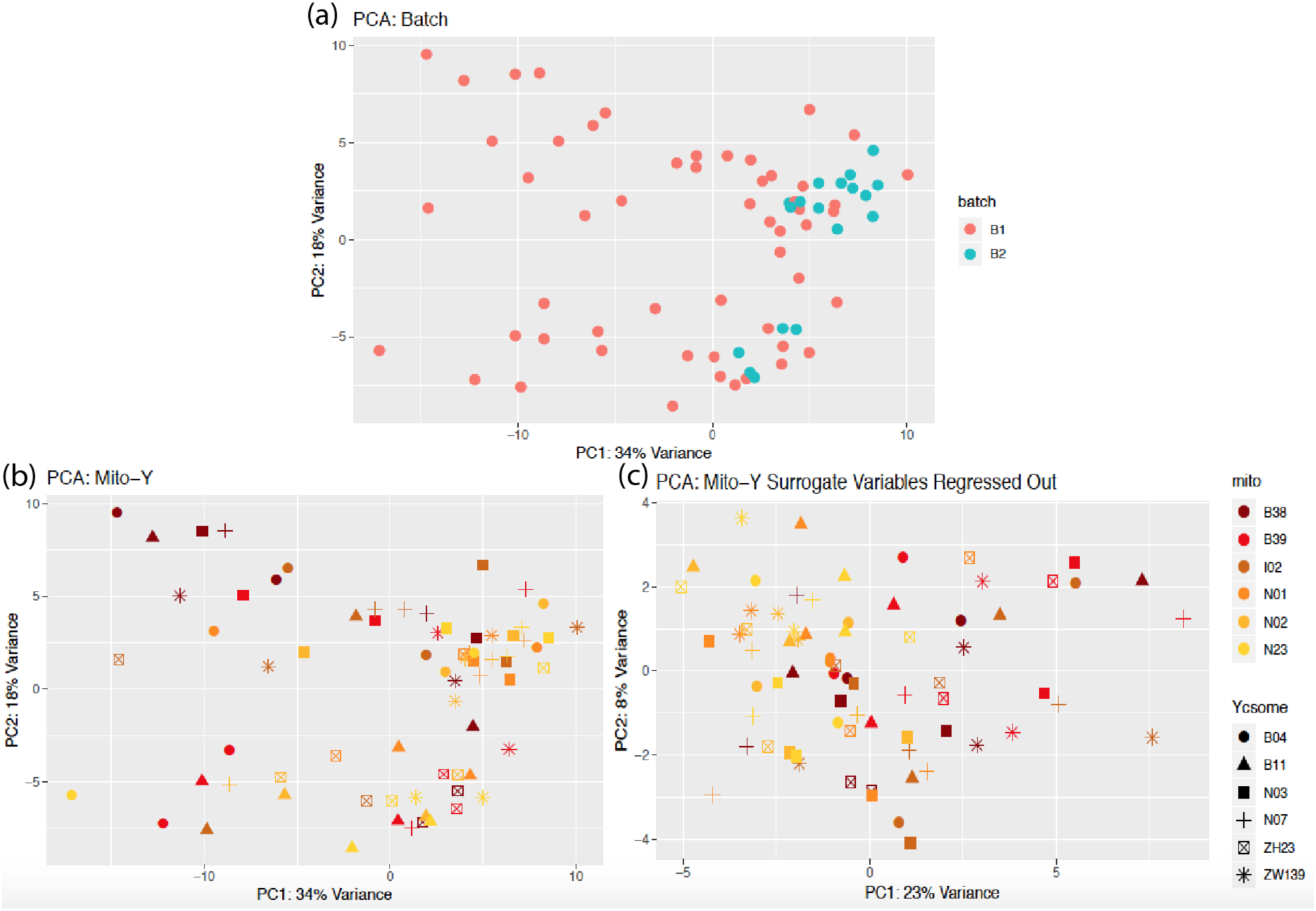
Principal components analysis on RNA-Seq samples. (a) Samples are coloured by the batch they were processed in and show a clear clustering of samples processed in the second batch (b) Samples are coloured by mitochondrial haplotype while the shape denotes Y haplotype, (c) Samples are plotted identically to (b) except that the surrogate variables have been regressed out of the data to show how their inclusion reduced the level of variation between samples.

### Surrogate Variable Analysis

Principal components analysis not only shows a clear batch effect amoungst our samples (Fig S2a) but also the failure of our samples to cluster by any measured grouping (i.e. mitochondrial haplogroup, Y haplotype, and occasionally replicate). This suggested the presence of unmeasured variables that may be impacting the sequencing data. To account for this, we used the R package *sva* which estimates the number of surrogate variables in the data. These variables can then be incorporated into all models to control for any unmeasured factors. The models used to test for differential expression across Y haplotype or mitochondrial haplogroup use the same full model with 8 surrogate variables identified by *sva* (∼ batch + surrogate_variable_1 +…+ surrogate_variable_8 + mitochondrial_haplogroup + Y_haplotype), while the test for differential expression due to mitochondrial-Y interactions uses a slightly different full model with an added interaction term and with 6 surrogate variables incorporated (∼ batch + surrogate_variable_1 +…+ surrogate_variable_6 + mitochondrial_haplogroup + Y_haplotype + mitochondrial_haplogroup:Y_haplotype).

**Figure S3.**
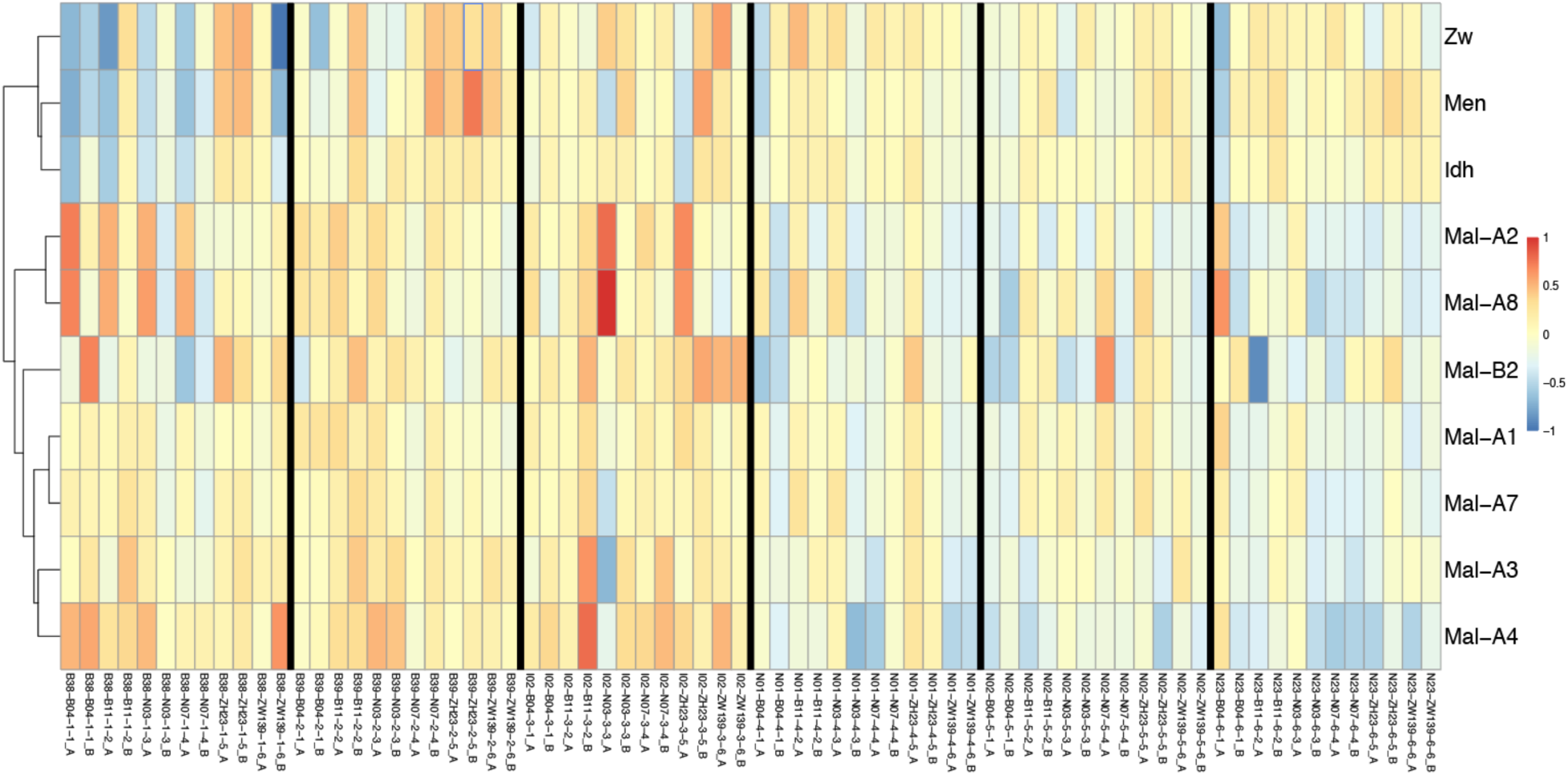
Expression of metabolic genes found to show significant variability in expression across mitochondrial haplogroups. Samples (36 mito-Y combinations with 2 replicates, labelled A and B, for each) and genes of interest are listed along the *x-* and *y-*axis respectively. Color indicates either overexpression (red) or under-expression (blue) compared to the mean expression level of that gene across all samples.

**Figure S4.**
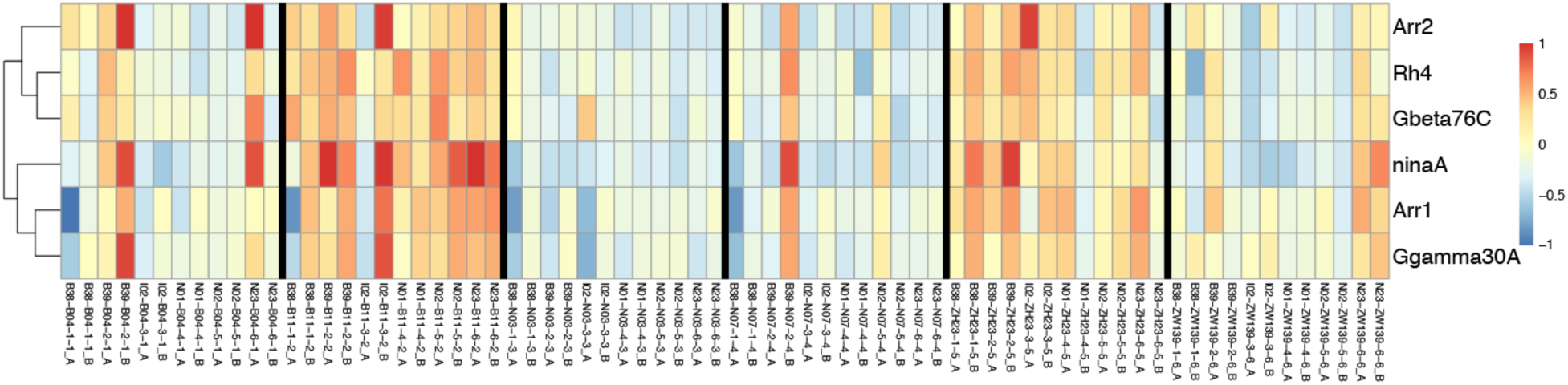
Expression of vision genes found to show significant variability in expression across Y haplotypes. Samples (36 mito-Y combinations with 2 replicates, labelled A and B, for each) and genes of interest are listed along the *x-* and *y-*axis respectively. Colour indicates either overexpression (red) or under-expression (blue) compared to the mean expression level of that gene across all samples.

**Figure S5.**
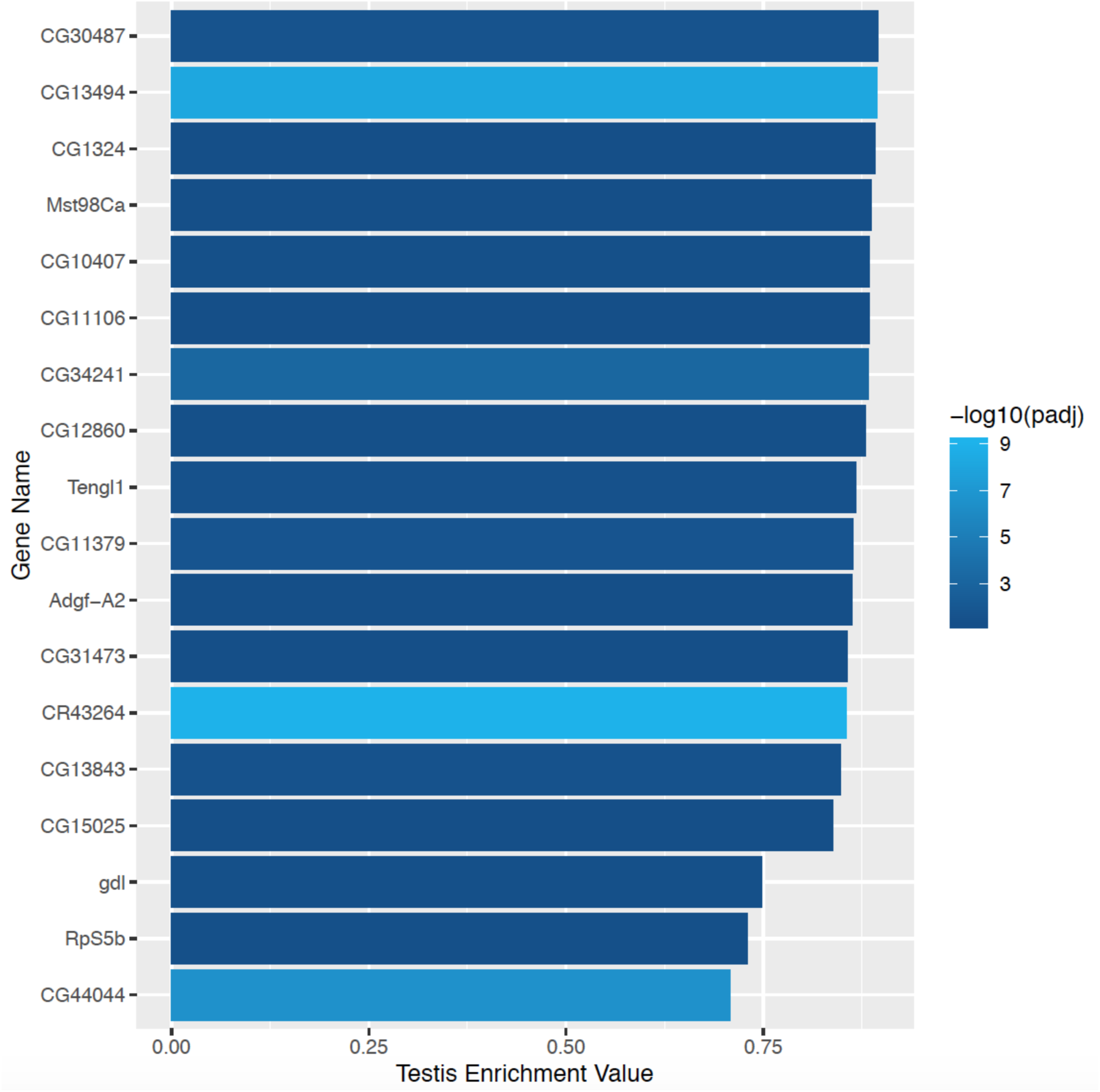
Y-Sensitive hits that are enriched for expression in the testis.

**Figure S6.**
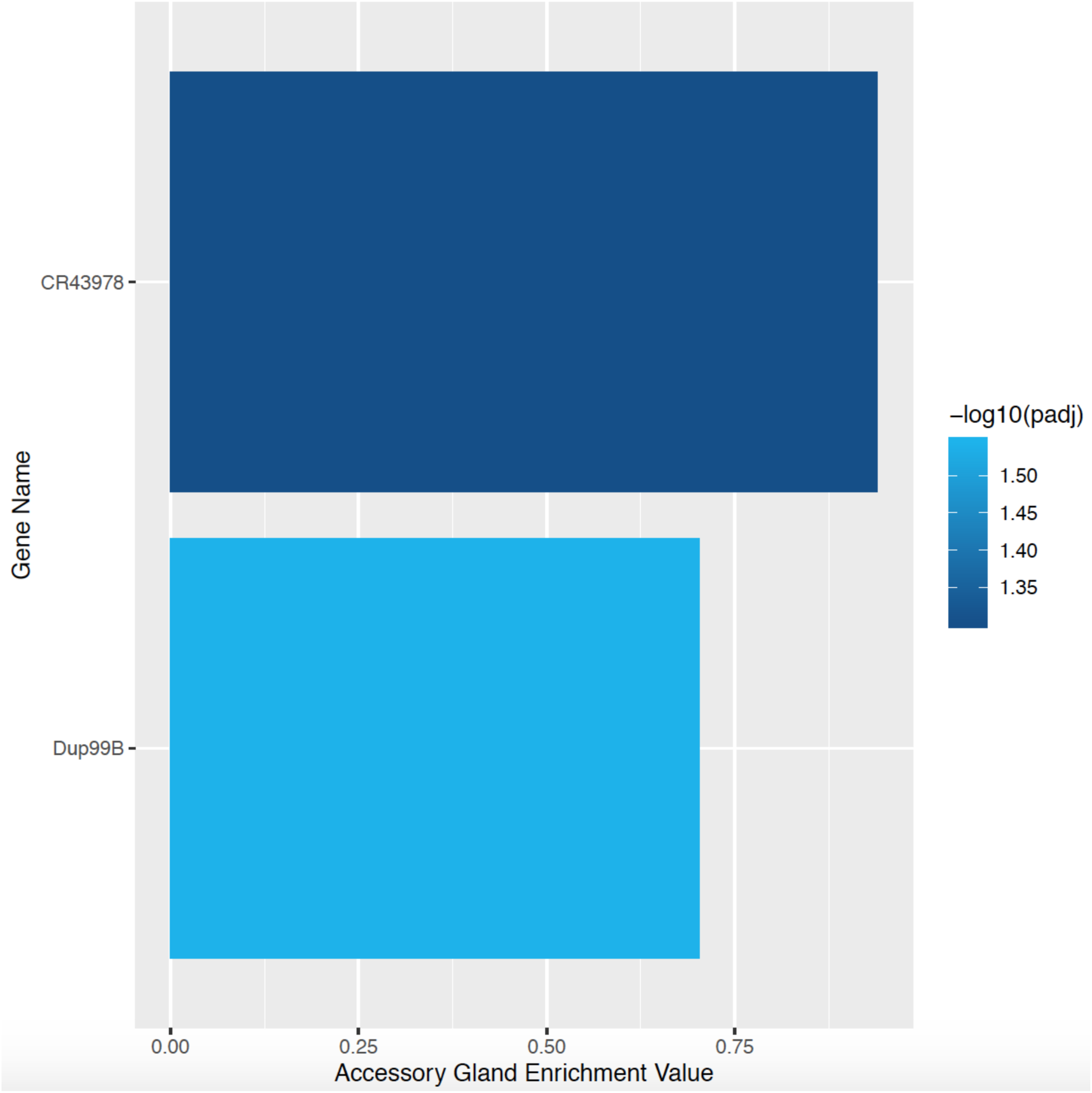
Y-Sensitive hits that are enriched for expression in the accessory glands.

**Figure S7.**
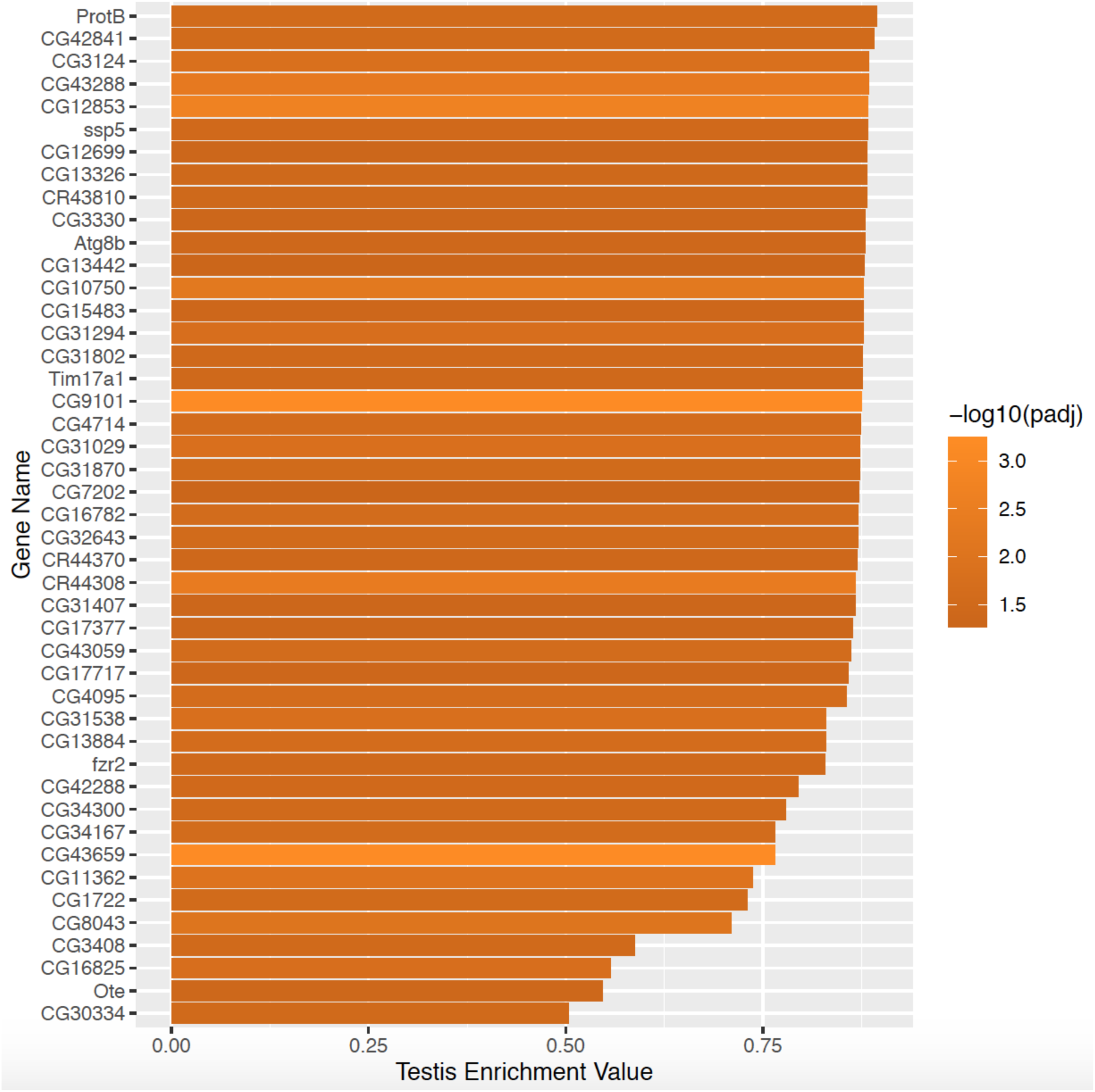
Mito-sensitive hits that are enriched for expression in the testis.

**Figure S8.**
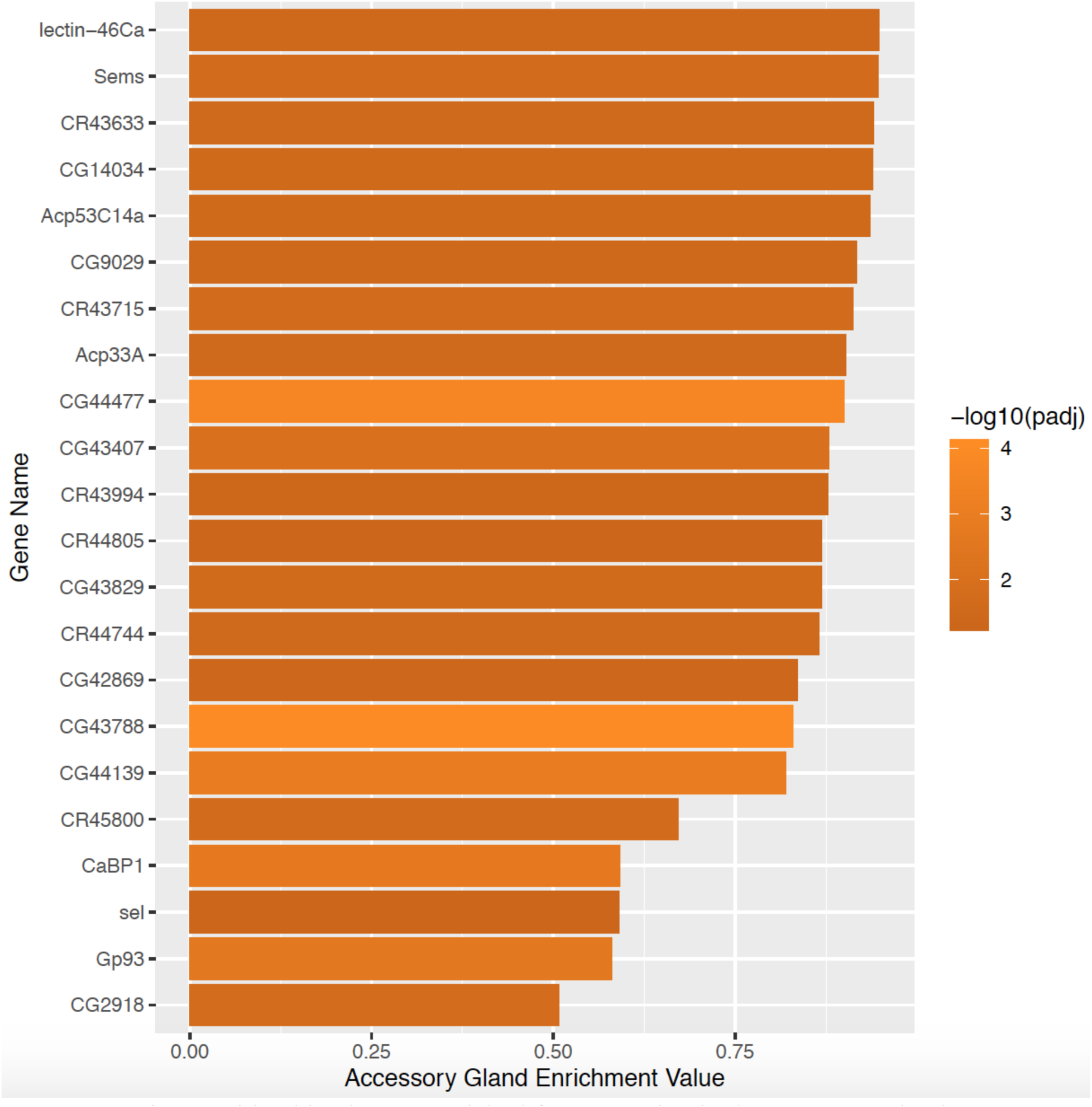
Mito-sensitive hits that are enriched for expression in the accessory glands.

## References

1. Gillham NW. 1994 Organelle Genes and Genomes. Oxford University Press.

2. Rand DM, Haney RA, Fry AJ. 2004 Cytonuclear coevolution: the genomics of cooperation. Trends Ecol. Evol. 19, 645–653.

3. Lane N. 2017 Origin of the eukaryotic cell. Mol. Front. J. 01, 108–120.

4. Blier PU, Dufresne F, Burton RS. 2001 Natural selection and the evolution of mtDNA-encoded peptides: evidence for intergenomic co-adaptation. Trends Genet. 17, 400–406.

5. Hill GE. 2015 Mitonuclear ecology. Mol. Biol. Evol. 32, 1917–1927.

6. Burton RS, Pereira RJ, Barreto FS. 2013 Cytonuclear genomic interactions and hybrid breakdown. Annu. Rev. Ecol. Evol. Syst. 44, 281–302.

7. Wade MJ, Goodnight CJ. 2006 Cyto-nuclear epistasis: two-locus random genetic drift in hermaphroditic and dioecious species. Evolution 60, 643–659.

8. Wade MJ, Drown DM. 2016 Nuclear-mitochondrial epistasis: a gene’s eye view of genomic conflict. Ecol. Evol. 6, 6460–6472.

9. Adams KL, Palmer JD. 2003 Evolution of mitochondrial gene content: gene loss and transfer to the nucleus. Mol. Phylogenet. Evol. 29, 380–395.

10. Berg OG, Kurland CG. 2000 Why mitochondrial genes are most often found in nuclei. Mol. Biol. Evol. 17, 951–961.

11. Lotz C et al. 2014 Characterization, design, and function of the mitochondrial proteome: from organs to organisms. J. Proteome Res. 13, 433–446.

12. Reinhardt K, Dowling DK, Morrow EH. 2013 Mitochondrial replacement, evolution, and the clinic. Science 341, 1345–1346.

13. Eyre-Walker A. 2017 Mitochondrial replacement therapy: Are mito-nuclear interactions likely to be a problem? Genetics 205, 1365–1372.

14. Birky CW Jr. 1995 Uniparental inheritance of mitochondrial and chloroplast genes: mechanisms and evolution. Proc. Natl. Acad. Sci. U. S. A. 92, 11331–11338.

15. Gemmell NJ, Metcalf VJ, Allendorf FW. 2004 Mother’s curse: the effect of mtDNA on individual fitness and population viability. Trends Ecol. Evol. 19, 238–244.

16. Frank SA, Hurst LD. 1996 Mitochondria and male disease. Nature 383, 224.

17. Charlesworth B, Charlesworth D. 1978 A model for the evolution of dioecy and gynodioecy. Am. Nat. 112, 975–997.

18. Frank SA. 1989 The evolutionary dynamics of cytoplasmic male sterility. Am. Nat. 133, 345–376.

19. Lewis D. 1941 Male sterility in natural populations of hermaphrodite plants the equilibrium between females and hermaphrodites to be expected with different types of inheritance. New Phytol. 46, 56–63.

20. Connallon T, Camus MF, Morrow EH, Dowling DK. 2018 Coadaptation of mitochondrial and nuclear genes, and the cost of mother’s curse. Proc. Biol. Sci. 285, 2257.

21. Budar F, Touzet P, De Paepe R. 2003 The nucleo-mitochondrial conflict in cytoplasmic male sterilities revisited. Genetica 117, 3–16.

22. Budar F, Pelletier G. 2001 Male sterility in plants: occurrence, determinism, significance and use. C. R. Acad. Sci. III 324, 543–550.

23. Kaul MLH. 1988 Male Sterility in Higher Plants. Springer.

24. Havird JC, Forsythe ES, Williams AM, Werren JH, Dowling DK, Sloan DB. 2019 Selfish mitonuclear conflict. Curr. Biol. 29, R496–R511.

25. Yee WKW, Rogell B, Lemos B, Dowling DK. 2015 Intergenomic interactions between mitochondrial and Y-linked genes shape male mating patterns and fertility in *Drosophila melanogaster*. Evolution 69, 2876–2890.

26. Dean R, Lemos B, Dowling DK. 2015 Context-dependent effects of Y chromosome and mitochondrial haplotype on male locomotive activity in *Drosophila melanogaster*. J. Evol. Biol. 28, 1861–1871.

27. Ågren JA, Munasinghe M, Clark AG. 2019 Sexual conflict through Mother’s Curse and Father’s Curse. Theor. Popul. Biol. 129, 9–17.

28. Lemos B, Branco AT, Hartl DL. 2010 Epigenetic effects of polymorphic Y chromosomes modulate chromatin components, immune response, and sexual conflict. Proc. Natl. Acad. Sci. U. S. A. 107, 15826–15831.

29. Lemos B, Araripe LO, Hartl DL. 2008 Polymorphic Y chromosomes harbor cryptic variation with manifold functional consequences. Science 319, 91–93.

30. Jiang P-P, Hartl DL, Lemos B. 2010 Y not a dead end: epistatic interactions between Y-linked regulatory polymorphisms and genetic background affect global gene expression in *Drosophila melanogaster*. Genetics 186, 109–118.

31. Kutch IC, Fedorka KM. 2017 A test for Y-linked additive and epistatic effects on surviving bacterial infections in *Drosophila melanogaster*. J. Evol. Biol. 30, 1400–1408.

32. Chippindale AK, Rice WR. 2001 Y chromosome polymorphism is a strong determinant of male fitness in *Drosophila melanogaster*. Proc. Natl. Acad. Sci. U. S. A. 98, 5677–5682.

33. Brown EJ, Bachtrog D. 2017 The Y chromosome contributes to sex-specific aging in Drosophila. bioRxiv 156042. (doi:10.1101/156042)

34. Innocenti P, Morrow EH, Dowling DK. 2011 Experimental evidence supports a sex-specific selective sieve in mitochondrial genome evolution. Science 332, 845–848.

35. Rand DM, Fry A, Sheldahl L. 2006 Nuclear-mitochondrial epistasis and drosophila aging: introgression of *Drosophila simulans* mtDNA modifies longevity in *D. melanogaster* nuclear backgrounds. Genetics 172, 329–341.

36. Zhu C-T, Ingelmo P, Rand DM. 2014 G×G×E for lifespan in Drosophila: mitochondrial, nuclear, and dietary interactions that modify longevity. PLoS Genet. 10, e1004354.

37. Clancy DJ. 2008 Variation in mitochondrial genotype has substantial lifespan effects which may be modulated by nuclear background. Aging Cell 7, 795–804.

38. Tower J. 2017 Sex-specific gene expression and life span regulation. Trends Endocrinol. Metab. 28, 735–747.

39. Grenier JK, Arguello JR, Moreira MC, Gottipati S, Mohammed J, Hackett SR, Boughton R, Greenberg AJ, Clark AG. 2015 Global diversity lines - a five-continent reference panel of sequenced *Drosophila melanogaster* strains. G3 (Bethesda) 5, 593–603.

40. Bolger AM, Lohse M, Usadel B. 2014 Trimmomatic: a flexible trimmer for Illumina sequence data. Bioinformatics 30, 2114–2120.

41. Hoskins RA et al. 2015 The Release 6 reference sequence of the *Drosophila melanogaster* genome. Genome Res. 25, 445–458.

42. Dobin A, Davis CA, Schlesinger F, Drenkow J, Zaleski C, Jha S, Batut P, Chaisson M, Gingeras TR. 2013 STAR: ultrafast universal RNA-seq aligner. Bioinformatics 29, 15–21.

43. Anders S, Pyl PT, Huber W. 2015 HTSeq—a Python framework to work with high-throughput sequencing data. Bioinformatics 31, 166–169.

44. Love MI, Huber W, Anders S. 2014 Moderated estimation of fold change and dispersion for RNA-seq data with DESeq2. Genome Biol. 15, 550.

45. Leek JT. 2014 svaseq: removing batch effects and other unwanted noise from sequencing data. Nucleic Acids Res. 42, e161.

46. Young MD, Wakefield MJ, Smyth GK, Oshlack A. 2010 Gene ontology analysis for RNA-seq: accounting for selection bias. Genome Biol. 11, R14.

47. Supek F, Bošnjak M, Škunca N, Šmuc T. 2011 REVIGO summarizes and visualizes long lists of gene ontology terms. PLoS One 6, e21800.

48. Leader DP, Krause SA, Pandit A, Davies SA, Dow JAT. 2018 FlyAtlas 2: a new version of the *Drosophila melanogaster* expression atlas with RNA-Seq, miRNA-Seq and sex-specific data. Nucleic Acids Res. 46, D809–D815.

49. Yoon JS, Gagen KP, Zhu DL. 1990 Longevity of 68 species of Drosophila. Ohio J. Sci. 90, 16–32.

50. Tower J, Arbeitman M. 2009 The genetics of gender and life span. J. Biol. 8, 38.

51. Camus MF, Clancy DJ, Dowling DK. 2012 Mitochondria, maternal inheritance, and male aging. Curr. Biol. 22, 1717–1721.

52. Ashburner M et al. 2000 Gene Ontology: tool for the unification of biology. Nat. Genet. 25, 25–29.

53. . The Gene Ontology Consortium. 2019 The Gene Ontology Resource: 20 years and still GOing strong. Nucleic Acids Res. 47, D330–D338.

54. Brosseau GE. 1960 Genetic analysis of the male fertility factors on the Y chromosome of *Drosophila melanogaster*. Genetics 45, 257–274.

55. Kennison JA. 1981 The genetic and cytological organization of the Y chromosome of *Drosophila melanogaster*. Genetics 98, 529–548.

56. Morgan TH. 1910 Sex limited inheritance in Drosophila. Science 32, 120–122.

57. Carvalho AB, Lazzaro BP, Clark AG. 2000 Y chromosomal fertility factors kl-2 and kl-3 of *Drosophila melanogaster* encode dynein heavy chain polypeptides. Proc. Natl. Acad. Sci. U. S. A. 97, 13239–13244.

58. Clark AG. 1990 Two tests of Y chromosomal variation in male fertility of *Drosophila melanogaster*. Genetics 125, 527–534.

59. Patel MR et al. 2016 A mitochondrial DNA hypomorph of cytochrome oxidase specifically impairs male fertility in *Drosophila melanogaster*. eLife 5.

60. Matsushita T, Fujii-Taira I, Tanaka Y, Homma KJ, Natori S. 2000 Male-specific IDGF, a novel gene encoding a membrane-bound extracellular signaling molecule expressed exclusively in testis of *Drosophila melanogaster*. J. Biol. Chem. 275, 36934–36941.

61. Schäfer M, Börsch D, Hülster A, Schäfer U. 1993 Expression of a gene duplication encoding conserved sperm tail proteins is translationally regulated in *Drosophila melanogaster*. Mol. Cell. Biol. 13, 1708–1718.

62. Ayyar S, Jiang J, Collu A, White-Cooper H, White RAH. 2003 Drosophila TGIF is essential for developmentally regulated transcription in spermatogenesis. Development 130, 2841–2852.

63. Jiang J, White-Cooper H. 2003 Transcriptional activation in Drosophila spermatogenesis involves the mutually dependent function of *aly* and a novel meiotic arrest gene *cookie monster*. Development 130, 563–573.

64. Saudan P et al. 2002 Ductus ejaculatorius peptide 99B (DUP99B), a novel *Drosophila melanogaster* sex-peptide pheromone. Eur. J. Biochem. 269, 989–997.

65. Rathke C, Barckmann B, Burkhard S, Jayaramaiah-Raja S, Roote J, Renkawitz-Pohl R. 2010 Distinct functions of Mst77F and protamines in nuclear shaping and chromatin condensation during Drosophila spermiogenesis. Eur. J. Cell Biol. 89, 326–338.

66. Raja SJ, Jayaramaiah Raja S, Renkawitz-Pohl R. 2006 Replacement by *Drosophila melanogaster* protamines and Mst77F of histones during chromatin condensation in late spermatids and role of sesame in the removal of these proteins from the male pronucleus. Mol. Cell. Biol. 26, 3682–3682.

67. Thurmond J et al. 2019 FlyBase 2.0: the next generation. Nucleic Acids Res. 47, D759– D765.

68. LaFlamme BA, Ravi Ram K, Wolfner MF. 2012 The *Drosophila melanogaster* seminal fluid protease ‘Seminase’ regulates proteolytic and post-mating reproductive processes. PLoS Genet. 8, e1002435.

69. Dowling DK, Tompkins DM, Gemmell NJ. 2015 The Trojan female technique for pest control: a candidate mitochondrial mutation confers low male fertility across diverse nuclear backgrounds in *Drosophila melanogaster*. Evol. Appl. 8, 871–880.

70. Yee WKW, Sutton KL, Dowling DK. 2013 In vivo male fertility is affected by naturally occurring mitochondrial haplotypes. Curr. Biol. 23, R55–6.

71. Clancy DJ, Hime GR, Shirras AD. 2011 Cytoplasmic male sterility in *Drosophila melanogaster* associated with a mitochondrial CYTB variant. Heredity 107, 374–376.

72. Yue L, Karr TL, Nathan DF, Swift H, Srinivasan S, Lindquist S. 1999 Genetic analysis of viable *Hsp90* alleles reveals a critical role in Drosophila spermatogenesis. Genetics 151, 1065–1079.

73. Wasbrough ER, Dorus S, Hester S, Howard-Murkin J, Lilley K, Wilkin E, Polpitiya A, Petritis K, Karr TL. 2010 The *Drosophila melanogaster* sperm proteome-II (DmSP-II). J. Proteomics 73, 2171–2185.

74. Cox DN, Chao A, Baker J, Chang L, Qiao D, Lin H. 1998 A novel class of evolutionarily conserved genes defined by piwi are essential for stem cell self-renewal. Genes Dev. 12, 3715–3727.

75. Lin H, Spradling AC. 1997 A novel group of *pumilio* mutations affects the asymmetric division of germline stem cells in the Drosophila ovary. Development 124, 2463–2476.

76. Malone CD, Brennecke J, Dus M, Stark A, McCombie WR, Sachidanandam R, Hannon GJ. 2009 Specialized piRNA pathways act in germline and somatic tissues of the Drosophila ovary. Cell 137, 522–535.

77. Huang X, Fejes Tóth K, Aravin AA. 2017 piRNA Biogenesis in *Drosophila melanogaster*. Trends Genet. 33, 882–894.

78. Tóth KF, Pezic D, Stuwe E, Webster A. 2016 The piRNA Pathway Guards the Germline Genome Against Transposable Elements. Adv. Exp. Med. Biol. 886, 51–77.

79. Vagin VV, Sigova A, Li C, Seitz H, Gvozdev V, Zamore PD. 2006 A distinct small RNA pathway silences selfish genetic elements in the germline. Science 313, 320–324.

80. Vagin VV et al. 2013 Minotaur is critical for primary piRNA biogenesis. RNA 19, 1064– 1077.

81. Gangaraju VK, Yin H, Weiner MM, Wang J, Huang XA, Lin H. 2011 Drosophila Piwi functions in Hsp90-mediated suppression of phenotypic variation. Nat. Genet. 43, 153–158.

82. Satoh AK, Ready DF. 2005 Arrestin1 mediates light-dependent rhodopsin endocytosis and cell survival. Curr. Biol. 15, 1722–1733.

83. Shieh B-H, Kristaponyte I, Hong Y. 2014 Distinct roles of arrestin 1 protein in photoreceptors during Drosophila development. J. Biol. Chem. 289, 18526–18534.

84. Katz B, Minke B. 2018 The Drosophila light-activated TRP and TRPL channels - Targets of the phosphoinositide signaling cascade. Prog. Retin. Eye Res. 66, 200–219.

85. Niemeyer BA, Suzuki E, Scott K, Jalink K, Zuker CS. 1996 The Drosophila light-activated conductance is composed of the two channels TRP and TRPL. Cell 85, 651–659.

86. Gu Y, Oberwinkler J, Postma M, Hardie RC. 2005 Mechanisms of light adaptation in Drosophila photoreceptors. Curr. Biol. 15, 1228–1234.

87. Dolph PJ, Man-Son-Hing H, Yarfitz S, Colley NJ, Deer JR, Spencer M, Hurley JB, Zuker CS. 1994 An eye-specific G beta subunit essential for termination of the phototransduction cascade. Nature 370, 59–61.

88. Hardie RC, Raghu P. 2001 Visual transduction in Drosophila. Nature 413, 186–193.

89. Hanlon CD, Andrew DJ. 2015 Outside-in signaling--a brief review of GPCR signaling with a focus on the Drosophila GPCR family. J. Cell Sci. 128, 3533–3542.

